# Immortalized Intestinal Telocytes - A Stem Cell Niche In Vitro

**DOI:** 10.1101/2024.11.27.625598

**Authors:** Marco Canella, Amal Gharbi, Deborah Duran, Noa Corem, Yinon Ben-Neriah, Liron Birimberg-Schwartz, Ittai Ben-Porath, Myriam Grunewald, Michal Shoshkes-Carmel

## Abstract

Telocytes, a distinct type of mesenchymal cell, have recently emerged as crucial components of the intestinal stem cell niche, primarily by providing essential Wnt proteins that drive stem cell function. However, their detailed biology and the specific mechanisms by which they regulate stem cell activity remain largely unexplored. Here, we established an immortalized intestinal telocyte cell line that retains the key cellular and molecular characteristics of native telocytes. Remarkably, this cell line, unlike telocyte-depleted immortalized mesenchyme, supports the growth of both mouse and human organoids in co-culture without the need for external factor supplementation. These findings indicate that telocytes are not only essential but are also sufficient to maintain stem cell activity. This immortalized telocyte line provides a valuable source of factors that support stem cell activity in vitro, offering a powerful tool for advancing our understanding of telocyte biology and the reciprocal communication between stem cells and their niche.

## Introduction

Telocytes, unique mesenchymal cells characterized by the expression of the transcription factor Foxl1, have recently been recognized as crucial for maintaining intestinal homeostasis^1–4^. Historically, telocytes were described primarily through electron microscopy as special interstitial cells present in the stroma of many organs^5–11^. They are distinct from fibroblasts due to their exceptionally long cellular extensions, known as telopodes, which can extend hundreds of microns from the cell body. These telopodes vary in width, and feature dilated regions called podoms, which contain mitochondria, caveolae, lipid rafts used for signal transduction, and elements of endoplasmic reticulum (ER).Through these extensions, telocytes form intricate 3D networks with other telocytes and diverse cell types, including epithelial cells, blood and lymphatic vessels, immune cells, fibroblasts, myofibroblasts and nerve bundles^8,10–12^. While they are present in many organs, the essential role of telocytes in maintaining homeostasis was first explored in the intestine^2^.

In the intestine, telocytes establish an extensive 3D network along the basal side of the crypt-villus epithelium, playing a critical role as the stem cell niche by supplying Wnt proteins essential for stem cell function. Additionally, telocytes facilitate enterocyte differentiation^3^. They serve as signaling hubs, expressing a wide array of signaling molecules from key pathways, including Wnt, Rspo, Bmp, Tgfβ and Shh. Notably, telocytes exhibit heterogeneity in their signaling molecule expression based on their position along the crypt-villus axis, correlating with active signaling pathways in the epithelium^2,3^.

The crucial role of telocytes in maintaining intestinal homeostasis highlights the need for effective strategies to establish in vitro sources of these cells while preserving their in vivo features and functions. This is essential for advancing our understanding of their biology and functional mechanisms. However, several challenges complicate the maintenance of primary telocyte cultures. Telocytes are present in very low numbers, constituting about 0.5% of the intestinal stromal population, as determined by FACS^2^. Their unique characteristics and long cellular extensions render them particularly vulnerable to damage during dissociation and sorting procedures. Although current protocols for isolating mesenchymal cells from mouse intestines can yield a relatively high proportion of viable telocytes^13^, their low abundance and non-proliferative nature present significant hurdles. Our goal was to develop a cell line system of telocytes for in vitro studies. Immortalization involves manipulating cells to bypass normal limits on cell division, allowing for ‘infinite’ proliferation in vitro. Immortalized cell lines provide a consistent supply of cells and facilitate genetic manipulations^14^. One widely used method for generating immortalized cell lines involves introducing viral oncogenes. Among these, infection with SV40 (simian vacuolating virus) proteins, particularly Large T-antigen^15^, is one of the most successful immortalizing genes. Immortalization by Large T-antigen is achieved through its (i) binding to the Retinoblastoma protein (Rb)-E2F complex, which relieves cell proliferation restrictions by activating of E2F-mediated transcription^16–20^, and (ii) binding and inhibition of p53 activity^21–30^. These mechanisms prevent growth arrest and senescence, resulting in continuous cell proliferation.

In this study, we utilized Foxl1Cre-driven reporter mice to isolate GFP^+^ telocytes from the mouse jejunum, and employed the SV40 Large T-antigen immortalization protocol to create an immortalized intestinal telocyte cell line. Our results demonstrate that this cell line retains key telocyte features, including the ability to form extensive networks via telopodes, cover large surface areas, and express factors known to be essential for supporting stem cell activity. Furthermore, the immortalized telocytes preserve molecular characteristics typical of native telocytes, showing high enrichment for PDGFRα, CD34, and Mcam. Remarkably, these immortalized cells support growth of organoids-derived from both mouse and human tissues, without requiring additional factor supplementation. We propose that this immortalized telocyte cell line can be used as a resource for niche factors to support stem cell activity in vitro. Moreover, this cell line represents a valuable tool for advancing our understanding of telocyte biology and their interactions with stem cells in health and disease.

## Results

### Immortalized telocytes retain structural characteristics

We developed an immortalized telocyte cell line by crossing Foxl1-promoter-driven Cre mice with Rosa-mTmG reporter mice, which express a plasma membrane-bound green fluorescence protein (GFP)^1^ (**Figure 1A** scheme). This approach allowed us to label telocytes with a membrane-tagged GFP (green), while Foxl1-negative cells were labeled with membrane-tagged tdTomato (red). We dissected the mouse jejunum and isolated the mesenchyme^13^, which was then cultured for two weeks to recover from potential dissociation damage. Following recovery, we dissociated the mesenchymal cells, FACS-sorted and plated GFP^+^ telocytes and tdTomato^+^ (telocyte-depleted) mesenchymal cells separately.

**Figure 1.**
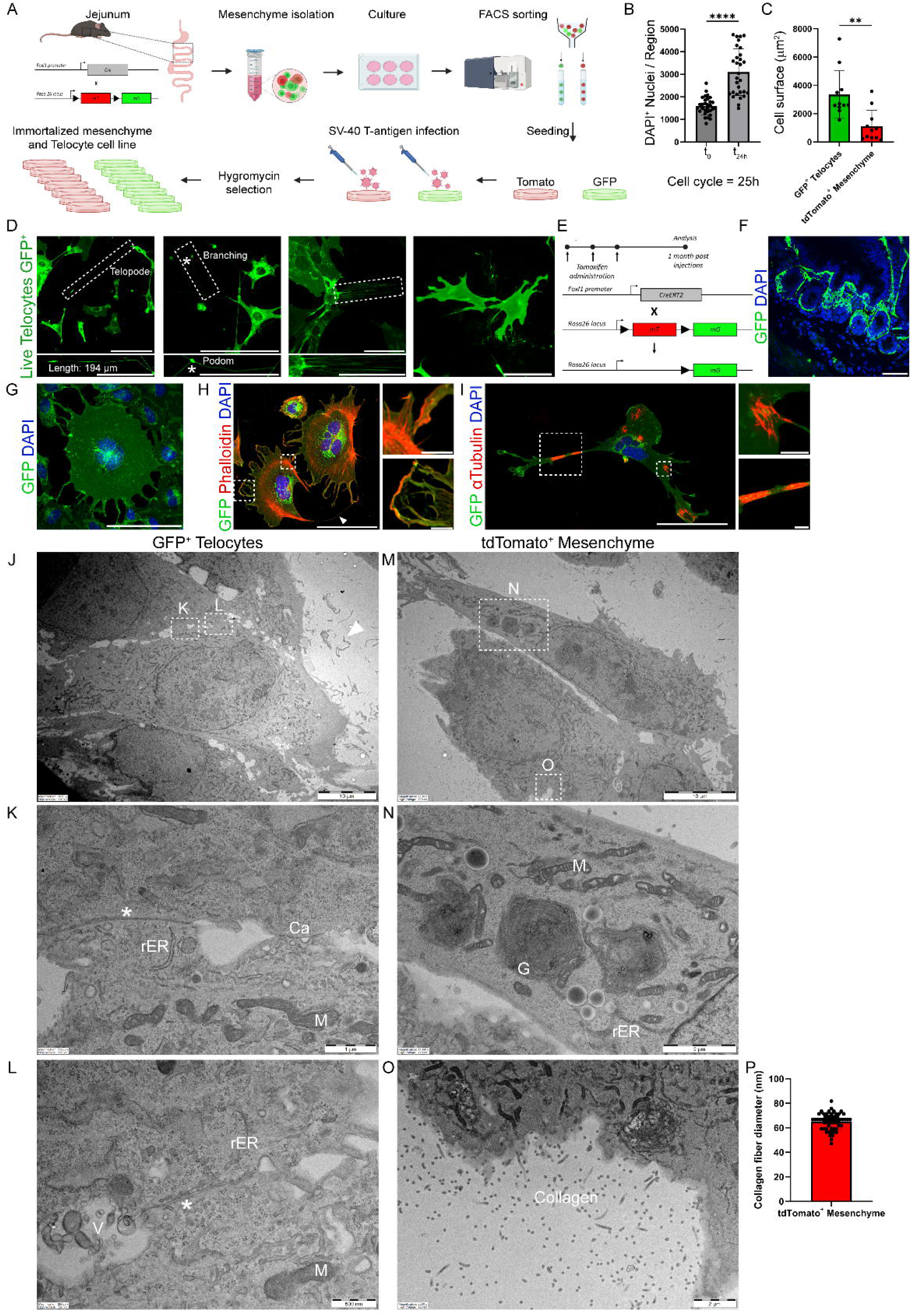
Immortalized telocytes retain structural characteristics. (**A**) Scheme illustrating the experimental strategy for creating the immortalized telocyte cell line. The jejunum from Foxl1-Cre; Rosa26-mTmG mouse model was dissected, and the mesenchyme was isolated and sorted into GFP^+^ telocytes and tdTomato^+^ (telocyte depleted) mesenchymal cells and cultured separately. Each cell population was then infected with SV40 T-antigen viral particles and selected with hygromycin to obtain the infected cells. (**B**) Quantification of DAPI^+^ nuclei at time 0 (t_0_) and after 24 hours (t_24h_) revealed a significant 94% increase in DAPI^+^ nuclei per region (n=30 regions per time point, p < 0.0001, Student’s t-test), suggesting a cell cycle duration of approximately 25 hours for immortalized telocyte cell line (**C**) Quantification of cell surface area revealed that GFP^+^ immortalized telocytes have a significantly larger surface compared to tdTomato^+^ (telocyte-depleted) immortalized mesenchymal cells (n=11 cells per group, <0.005, Student’s t-test). (**D**) Live imaging of immortalized telocyte cell culture revealed key cellular characteristics of these cells. The boxed regions highlight the length of telopodes (194 µm), their branching pattern, and the presence of dilated regions, known as podoms (indicated by asterisks). Additionally, immortalized telocytes form a network by interconnecting through several telopodes and cellular protrusions. (**E**) Schematic representation of the induction strategy in the Foxl1-CreERT2; Rosa26-mTmG mouse model. Tamoxifen was administered for three consecutive days, and the mouse was sacrificed after a month for jejunum analysis. (**F**) Immunofluorescence of the Foxl1-CreERT2; Rosa26-mTmG jejunum, stained for GFP (green) and DAPI (blue), revealed an intricate network of thin and flat Foxl1^+^ GFP^+^ telocytes forming a continuous subepithelial network lining the crypt-villus epithelium. Scale bar 50 µm. (**G**) Immunofluorescence staining of immortalized telocytes for GFP (green) and DAPI (blue) revealed their thin, flat morphology and demonstrated that their interconnected network is preserved in culture. (**H**) Immunofluorescence of immortalized telocytes stained for phalloidin (red) revealed asymmetrical distribution of actin filaments beneath the cell membrane and along cellular extensions. (**I**) Immunofluorescence of immortalized telocytes stained for α-tubulin (red) revealed the distribution of both thick and thin microtubule fibers, including branched structures across various cell compartments. Scale bar 100 µm, 10 µm. (**J–O**) Transmission Electron Microscopy images of GFP+ immortalized telocytes (J-L) and tdTomato+ (telocyte-depleted) immortalized mesenchymal cells (M-O) revealed unique ultrastructural characteristics of telocytes compared to fibroblasts. Telocyte-depleted mesenchymal cells often featured prominent Golgi apparatus (N) which was challenging to observe in immortalized telocytes. Cell-cell contact was a striking feature of immortalized telocytes, distinguishing them from telocyte-depleted mesenchymal cells. Caveolae, were predominantly clustered along broad, planar telocye-telocyte contact sites (K). Vesicle aggregation was frequently observed in the extracellular vicinity of telocytes (L). The cytoplasm of telocytes was densely populated with mitochondria and rough endoplasmic reticulum (rER), the latter being fully occupied by ribosomes. Surrounding the telocyte-depleted mesenchymal cells, abundant collagen fibers were present (O). (P) Quantification of the diameter of collagen fibers in tdTomato+ (telocyte-depleted) mesenchymal cell surrounding (n=60, 65.2 ±6.5). G - Golgi apparatus, rER - rough Endoplasmic Reticulum, Ca - Caveolae, M - Mitochondria, V - Vesicle aggregation. (asterisk point on planar telocyte-telocyte contact).

To achieve immortalization, we employed the Simian Virus 40 (SV40) Large T-antigen protocol, a well-established method known for its simplicity and reliability in immortalizing various cell types^31–33^. We infected GFP^+^ telocytes and tdTomato^+^ (telocyte-depleted) mesenchymal cells with viral particles expressing the SV40 Large T-antigen. The viral particles were produced by transfecting a lentiviral vector encoding the protein into HEK293T cells. Immortalized cells were expanded through 7-9 passages before being frozen for further analysis.

The term “immortalized” refers to cells with significantly higher proliferative capacity compared to native telocytes. Native telocytes exhibit exceptionally low proliferation rate, as demonstrated in the Grem1-driven Cre reporter mouse line^34^, where, over the course of a year, a single Grem1+ cell was observed to gradually expand and repopulate the entire subepithelial telocyte network^34^. In contrast, the immortalized telocyte cell line displayed a cell cycle duration of 25 hours, as determined by quantifying DAPI+ nuclei over a 24-hour culture period (**Figure 1B**).

The cell line was established two years ago, and the cells used in this manuscript were at a maximum of passage 18. Notably, these cells consistently retained the key features observed in earlier passages of the cell line.

Immortalization can alter cellular phenotypes; therefore, we assessed the characteristics of our immortalized telocyte cell line. The immortalized telocyte cells demonstrated high proliferation rates and variable levels of GFP expression. Morphologically, the cells appeared flat and thin, with projections that either extended hundreds of microns from the cell body (**Figure 1D**) or remained short and compact (**Figure 1G**). Dilated regions along these projections displayed high GFP signals, likely corresponding to the podom regions of the telopodes (**Figure 1D** boxed regions). During proliferation, the cells adopted a condensed and rounded morphology, extending multiple cellular protrusions. Following division, they spread out into thin, flat structures covering a large surface area (**Video 1** and **Figure 1G**), resembling the morphology observed along the intestinal crypt-villus epithelium (**Figures 1E-F**).

Compared to tdTomato+ (telocyte-depleted) mesenchyme (**Video 2**), GFP+ telocytes were at least three times larger (**Figure 1C**), exhibited highly dynamic cellular processes, and spread into thin amorphous structures that approached and made close contact with neighboring cells. Notably, dividing cells maintained intercellular contact and extended protrusions, forming interconnected networks (**Video 1)**. The cells were either mono- or multi-nucleated, a phenomenon whose origin remains unclear - whether it represents an artifact of immortalization or an intermediate state not typically observed in vivo.

The cells displayed pronounced polarization, with cytoskeletal elements concentrated in specific regions. Actin filaments, stained with phalloidin, were distributed asymmetrically beneath the plasma membrane (**Figure 1H**) and arranged in a regular striped pattern, suggesting contractility^35,36^. Cellular extensions were rich in actin, featuring bundles of parallel filaments, lamellipodia or membrane-ruffles (**Figure 1H** boxed regions), sometimes continuous with stress fibers (**Figure 1H** arrowhead). Microtubule distribution, stained for α-tubulin (**Figure 1I)** revealed a distinct pattern: thick, wide bundles along broader extensions, and thin, branched fibers at the base of thinner extensions and specific sites within the cytoplasm (**Figure 1I** boxed regions). This suggests that thick bundles of microtubules may represent stable structures, while thin, branched fibers indicate dynamic microtubules^37,38^.

Telocytes are primarily identified by electron microscopy, characterized by their unique ultrastructural features and their extremely long, thin telopodes. To analyze these features, we used transmission electron microscopy (TEM) to study the ultrastructure of both immortalized telocytes and telocyte-depleted mesenchyme. However, due to the close proximity of cells cultured in monolayer, we were unable to fully characterize the telopodes. In this setting, the telopodes appeared shorter and thinner and were often excluded from the 80nm ultrathin sections. Nonetheless, we observed portions of membranous extensions in the surrounding regions of the cells (**Figure 1J** arrowhead). Despite these limitations, distinct ultrastructural differences were identified between immortalized telocytes and telocyte-depleted mesenchyme. The nuclei of immortalized telocytes exhibited evenly distributed electron-dense heterochromatin puncta, whereas the nuclei of telocyte-depleted mesenchymal cells displayed heterochromatin in a more patch-like distribution. Additionally, telocyte-depleted mesenchymal cells often featured prominent Golgi apparatus (**Figure 1N**), while the Golgi complex was challenging to observe in immortalized telocytes—a characteristic consistent with native telocytes compared to fibroblasts^12^.Cell-cell contact was a striking feature of immortalized telocytes, distinguishing them from telocyte-depleted mesenchymal cells, which exhibited significantly less interaction even when in close proximity (**Figures 1J** and **1M, Videos 1** and **2**). Caveolae (**Figure 1K**, Ca), small invaginations of the plasma membrane typically enriched in proteins and lipids involved in signal transduction and a hallmark of telocytes^11,39^, were predominantly clustered along broad, planar telocye-telocyte contact sites (**Figures 1K** and **1L**, asterisks). Vesicle aggregation, another characteristic feature of telocytes^5,12^, with diameters ranging from 50-200nm, was frequently observed in the extracellular vicinity (**Figure 1L, V**).

The cytoplasm of telocytes was densely populated with mitochondria and rough endoplasmic reticulum (rER), the latter being fully occupied by ribosomes. The distribution of rER and mitochondria was less organized and less partitioned in telocytes compared to telocyte-depleted mesenchymal cells. Surrounding the telocyte-depleted mesenchymal cells, abundant collagen fibers with an average diameter of 60 nm were observed (**Figure 1O** and **1P**), consistent with the primary role of fibroblasts in extracellular matrix production. Notably, collagen fibers were absent in cultured telocytes, highlighting the distinct identity of telocytes compared to fibroblasts.

Thus, immortalized telocytes preserve key morphological and ultrastructural features of native telocytes.

### Immortalized telocytes preserve molecular expression profile

In vivo, intestinal telocytes are a distinct mesenchymal population, characterized by their expression of PDGFRα, CD34 and Foxl1. The diverse expression of signaling molecules in these cells highlights the heterogeneity within this population, which likely arises from complex interactions with surrounding cells. However, it was unclear whether telocytes that are immortalized and cultured outside their native environment, isolated from other cell types, would retain their molecular markers. To evaluate the molecular profile of these immortalized cells, we first validated Foxl1 expression, which drives our reporter mice, using single-molecule RNA fluorescence in situ hybridization (smFISH). Our analysis confirmed that Foxl1 mRNA is expressed in GFP^+^ telocytes, while its expression remains low in the tdTomato^+^ mesenchymal cells lacking telocytes (**Figures 2A-C**). This result confirms that, despite immortalization, the cell populations can still be distinguished by their Foxl1 expression.

**Figure 2.**
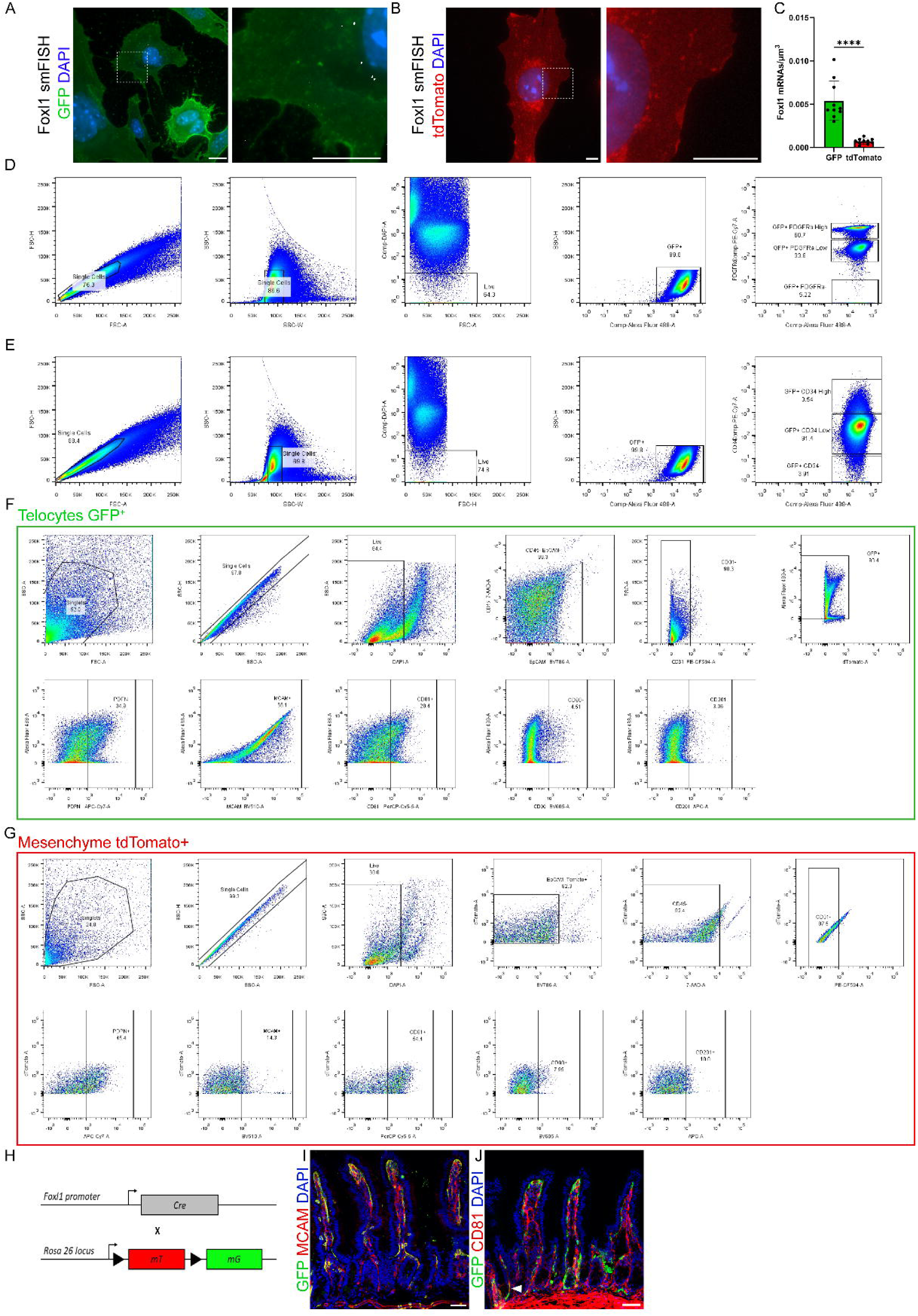
Immortalized telocytes preserve molecular expression profile. (**A–B**) Single-molecule RNA in situ fluorescence hybridization (smFISH) for Foxl1 mRNA (white) shows expression in GFP^+^ immortalized telocytes while minimal Foxl1 expression is detected in tdTomato^+^ (telocyte-depleted) immortalized mesenchymal cells. Scale bar 10 µm. (**C**) Quantification of Foxl1 mRNA molecules in GFP+ immortalized telocytes versus tdTomato+ mesenchymal cells (n=10 images per group, p<0.0001, Student’s t-test). (**D–E**) FACS analysis of GFP^+^ immortalized telocytes stained for PDGFRα (D) or CD34 (E) reveals two populations: PDGFRα^High^ and PDGFRα^Low^, with the majority of GFP^+^ immortalized telocytes are CD34^Low^. (**F–G**) FACS analysis comparing GFP+ immortalized telocytes to tdTomato^+^ mesenchymal cells for marker expression shows that GFP^+^ telocytes are enriched for Mcam expression compared to tdTomato^+^ mesenchymal cells, whereas Cd81 is more abundant in tdTomato^+^ mesenchymal cells. (**H-J**) Immunofluorescence staining of jejunum sections from Foxl1Cre; Rosa-mTmG mice shows that Mcam is predominantly expressed in GFP+ telocytes (I). Cd81 is expressed in GFP+ telocytes located along the crypt epithelium (J, arrowhead).

Next, we assessed two additional molecular hallmarks of intestinal telocytes, PDGFRα and CD34, using FACS analysis and immunofluorescence staining (**Figures 2D-G**). The majority of the immortalized telocytes expressed both PDGFRα and CD34. We observed two distinct populations based on PDGFRα expression levels: a PDGFRα^high^ population and a PDGFRα ^low^ population (**Figures 2D-E**).

To further characterize the differential expression profiles between GFP^+^ telocytes and tdTomato^+^ (telocyte-depleted) mesenchymal cells, we conducted FACS analysis using additional mesenchymal markers. We included markers of epithelial cells (EpCAM^+^), immune cells (CD45^+^) and endothelial cells (CD31^+^) to confirm the culture’s purity. We then stained for various mesenchymal markers, including the mucin-type glycoprotein Podoplanin (Pdpn), melanoma cell adhesion protein (Mcam), and the cell surface glycoproteins CD81, CD90 (Thy1) and CD201 (**Figures 2F-G**). Our analysis revealed significant differences in marker expression: Mcam was found in over 65% of GFP^+^ telocytes compared to just 14% of tdTomato^+^ mesenchymal cells, while CD81 was present in 28% of GFP^+^ cells but 54% of tdTomato^+^ cells. To examine the expression patterns of these markers and their relationship to native intestinal telocytes, we performed immunofluorescence staining on jejunum sections from Foxl1Cre; Rosa-mTmG mice, where telocytes are labeled with membrane-bound GFP (green) (**Figures 2H-J**). We found that Mcam was co-expressed in the majority of GFP^+^ telocytes (**Figure 2I**). In contrast, CD81 showed minimal co-localization, being restricted to a subset of telocytes located along the crypt epithelium (**Figure 2J** arrowhead).

Similarly, immunofluorescence staining of immortalized telocytes showed that all GFP+ telocytes consistently expressed PDGFRα (**Figure S1A**), while subsets expressed CD34 (**Figure S1B**), and others co-expressed αSMA and Thy1. Importantly, these expression profiles closely align with those observed in native telocytes (Gharbi et al., work in progress), reinforcing the validity of the immortalized telocyte model.

These findings demonstrate that immortalized telocytes maintain the original molecular profile of a heterogenous mesenchymal population, showing high enrichment for Foxl1, PDGFRα, CD34 and Mcam expression, effectively preserving their key characteristics and markers.

### Immortalized telocytes exhibit matrix remodeling and contraction skills

We next aimed to explore functional characteristics of telocytes that are difficult to study in vivo but can be effectively examined using in vitro cultures. Given the polarized distribution of cytoskeletal elements in GFP^+^ immortalized telocytes, we focused on their potential to contract and remodel the extracellular matrix (ECM). We stained GFP^+^ immortalized telocytes for matrix metalloproteinase 10 (Mmp10), known for its role in ECM degradation and clinical relevance in modulating inflammatory bowel disease (IBD)^40^ and a-smooth muscle actin (aSMA), a marker linked to contractile activity^41^. Both Mmp10 and aSMA were positively stained in a subset of the GFP^+^ immortalized telocytes (**Figure 3A**), highlighting that even when detached from their native environment, these immortalized telocytes retain significant heterogeneity.

**Figure 3.**
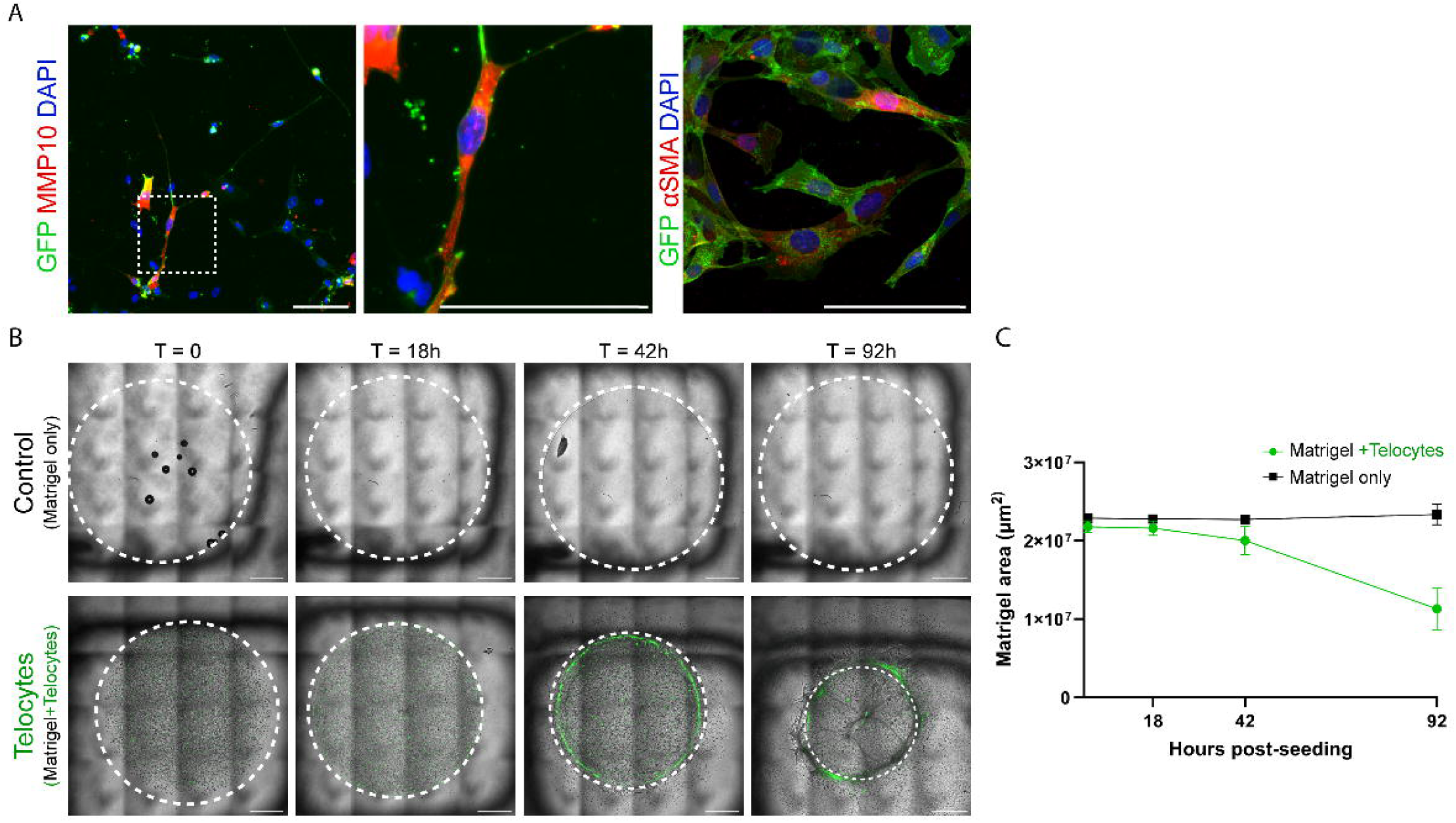
Immortalized telocytes exhibit matrix remodeling and contraction skills. **(A**) Immunofluorescence of GFP^+^ immortalized telocytes for matrix metalloproteinase10 and alpha smooth muscle actin (αSMA) (red) reveals expression in a subset of GFP^+^ immortalized telocytes (green). (**B**) Confocal images of Matrigel domes with and without 10,000 GFP^+^ immortalized telocytes. Images were acquired 0, 18, 42 and 92hours post-seeding. Scale bar 1 mm. (**C**) Quantification of Matrigel area with and without GFP^+^ immortalized telocytes. (n=4 biological replicates).

Next, we utilized Matrigel-based contraction assay to evaluate the telocytes’ ECM remodeling capabilities. This assay measures the contraction of the Matrigel dome to reflect cell-extracellular matrix interactions and cellular 3D contractility. Immortalized telocytes were embedded in Matrigel at a density of 10,000 cells per 30μl of Matrigel. Forty-two hours after seeding, the telocytes extended their processes and spread across the Matrigel dome (**Figures 3B-C**). We observed a significant contraction of the gel, indicating the telocytes’ active participation in ECM manipulation. This contraction, along with the morphological rearrangement of telocytes, suggests their potential to influence tissue architecture through mechanical forces, underscoring their role in the matrix environment.

### Immortalized telocytes retain expression of stem cell niche signaling molecules

In vivo, telocytes play a crucial role as a signaling hub maintaining intestinal homeostasis by supporting both stem cell function and enterocyte differentiation^1,2^. To determine if immortalized telocytes retain these niche functions, we assessed their expression of key signaling molecules.

Immunofluorescence staining revealed that a subset of immortalized telocytes expressed Porcupine (Porcn), an O-acyltrasferase localized in the endoplasmic reticulum, crucial for Wnt secretion (**Figure 4A**). This observation suggests that telocyte may exhibit heterogeneity in their ability to secrete Wnt proteins. Additionally, Vangl2, a receptor involved in the non-canonical Wnt-signaling pathway, displayed variable expression levels and a polarized distribution along the telocytes’ extensions (**Figure 4B**). This pattern suggests that telocytes not only support signaling but also have the capability to sense and respond to these signals.

**Figure 4.**
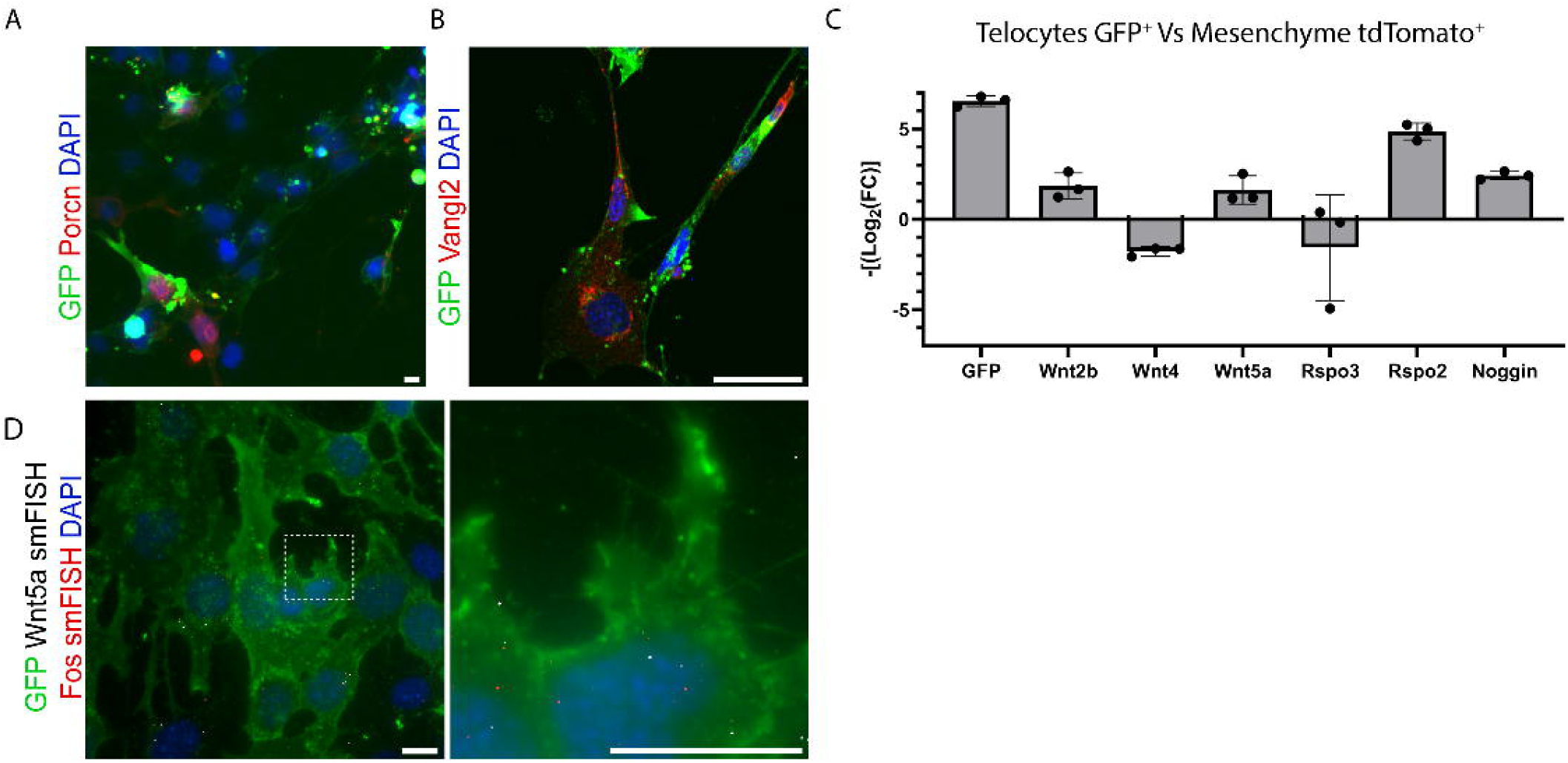
Immortalized telocytes retain expression of stem cell niche signaling molecules. (**A–B**) Immunofluorescence of GFP^+^ immortalized telocytes for the Wnt modifier enzyme, Porcupine and the non-canonical Wnt receptor Vangolin2 (red) reveals their expression in a subset of GFP^+^ immortalized telocytes. Scale bar 100 µm. (**C**) RT-qPCR analysis of GFP^+^ immortalized telocytes reveals the expression of stem cell niche signaling molecules in GFP^+^ immortalized telocytes. Data from three experiments (n=3). Each result is represented as an individual dot, and error bars indicate the standard deviation. (**D**) smFISH for Wnt5a (white) and Fos (red) reveals the expression of the signaling molecules in GFP^+^ immortalized telocytes. Scale bar 10 µm, 1 µm.

Quantitative PCR analysis further confirmed that immortalized telocytes retain key functions of the stem cell niche. Compared to mesenchymal telocyte-depleted population, immortalized telocytes showed significant enrichment of the canonical Wnt2b ligand (**Figure 4C).** Wnt2b, which is expressed in native telocytes along the crypt epithelium^2^, is known to regulate intestinal inflammatory cytokines^42^, and mutations in this gene are linked to neonatal-onset chronic diarrhea in humans^43^.

In addition to Wnt2b, immortalized telocytes were enriched for the non-canonical Wnt5a, a ligand highly expressed in native villus-tip telocytes^3^. This enrichment further indicates that our cell line encompasses diverse populations of telocytes. Furthermore, Rspo2, a crucial Wnt inducer, and Noggin, a Bmp inhibitor - both vital for the stem cell niche - were also enriched in telocytes compared to mesenchymal telocyte-depleted population.

Interestingly, although Rspo3 was expressed in telocytes, its expression was higher in the telocyte-depleted population compared to telocytes, reflecting the in vivo situation^2^. smFISH for Wnt5a validated the presence of mRNA molecules within telocytes, showing close proximity to the expression of mRNAs of the AP-1 transcription factor component Fos (**Figure 4D)**. This finding further supports the role of teloytes in both supporting and responding to signaling cues. Thus, immortalized telocytes maintain their heterogeneity, encompassing diverse populations of telocytes that express stem cell niche factors in vitro.

### Immortalized telocytes support organoid growth without additional growth factor supplements

Finally, we investigated whether immortalized telocytes retain their ability to maintain stem cell niche activity in vitro. Stem cells are characterized by their capacity to self-renew and differentiate into specialized progeny. Organoids - 3D tissue models formed by the proliferation and differentiation of either pluripotent or adult stem cells within a matrix scaffold - serve as a powerful tool in both fundamental and biomedical research^44–47^. These models have been successfully established for various organs and tissues in both mice and humans.

The generation of organoids from adult or induced pluripotent human stem cells has opened up opportunities for modeling human development, disease, drug discovery and personalized medicine ^44,48^. In these models, stem cell-containing crypts are isolated and embedded in Matrigel as an ex vivo replacement for the ECM. Crucially, the establishment and maintenance of organoids require supplementation with niche factors that support the formation of organized and functional tissue^44–48^.

To assess the potential of immortalized telocytes to support organoid growth without additional niche factors, we co-cultured these cells with mouse small intestinal-derived organoids embedded in Matrigel. This setup was compared with tdTomato^+^ (telocyte-depleted) mesenchymal cells, as well as organoids supplemented with or without external niche factors: EGF, Noggin, Rspondin and FGF (ENRF) (**Figure 5A** experimental design). By day 1 of culture, organoids in co-culture with GFP^+^ immortalized telocytes exhibited the largest sphere morphology compared to all other conditions (**Figures 5B-C**).

**Figure 5.**
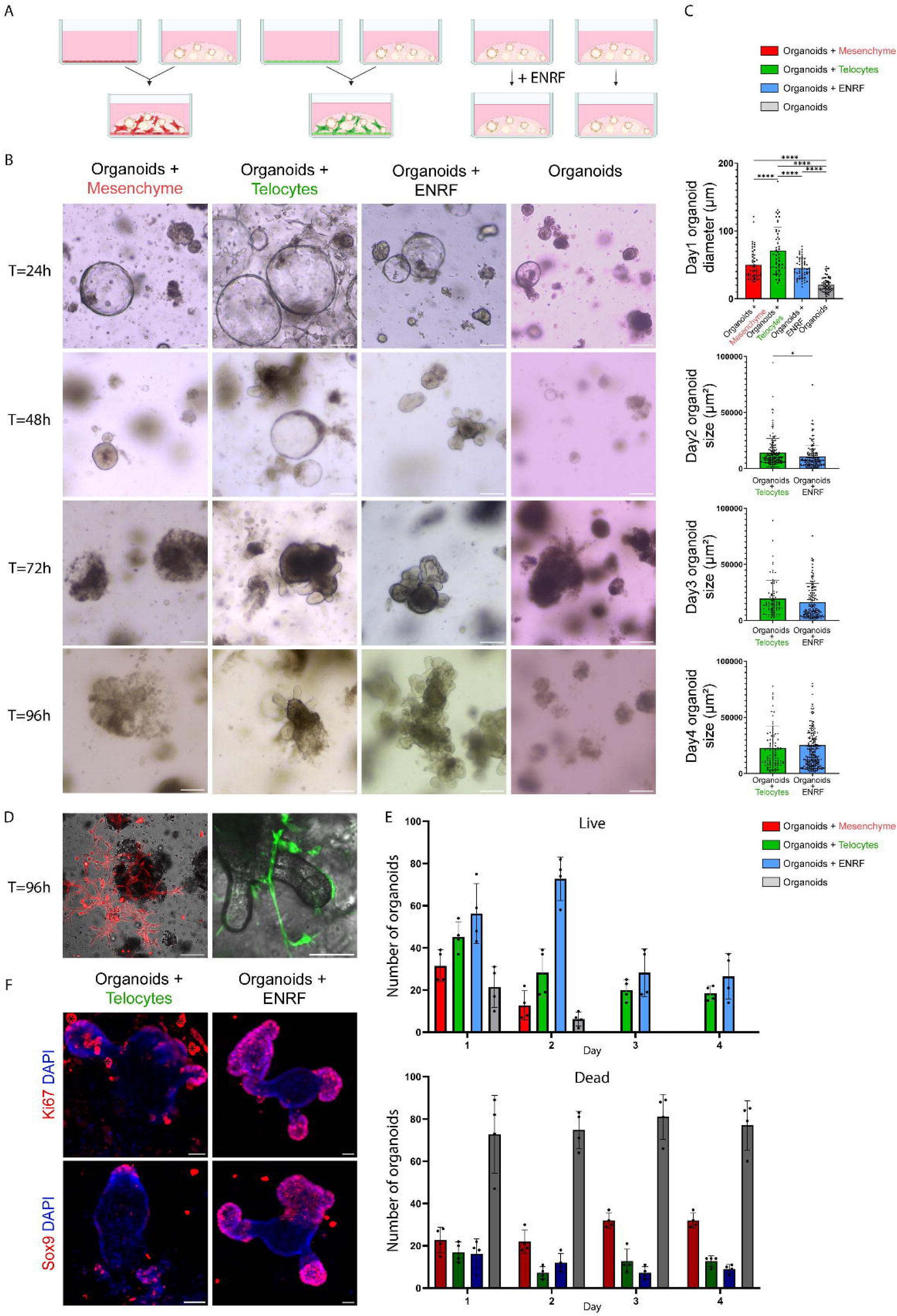
Immortalized telocytes support organoid growth without additional growth factor supplements. (**A**) Schematic illustration of organoid co-culture strategy. Mouse small intestinal organoids were cultured under four conditions: in co-cultured with tdTomato+ immortalized mesenchymal cells, in co-culture with GFP^+^ immortalized telocytes, supplemented with Egf, Noggin, Rspo1 and FGF or organoids only (**B**) Photographs of cultured organoids, taken daily over four days, show that organoids die within 24 hours under all conditions except when co-cultured with GFP^+^ immortalized telocytes or supplemented with external factors. (**C**) Quantification of organoid diameters 24 hours after seeding shows that organoids co-cultured with GFP+ immortalized telocytes formed significantly larger spheroids compared to all other conditions (n=4 replicates, at least 50 organoids per group, p< 0.005, one-way ANOVA with multiple comparisons). By day 2, organoids co-cultured with GFP+ immortalized telocytes were significantly larger than those cultured with growth factors alone (n=4 replicates, at least 50 organoids per group, p < 0.05, Student’s t-test). At days 3 and 4, no significant differences in organoid size were observed between co-cultures with telocytes and those supplemented with growth factors. (**D**) Confocal imaging shows GFP+ immortalized telocytes establishing close contact with organoids, whereas tdTomato+ immortalized mesenchymal cells are distributed within the Matrigel alongside dying organoids. (**E**) Quantification of live and dead organoids over four days of culture (n=4 replicates, at least 50 organoids per group) reveals that by day 4, only organoids co-cultured with telocytes or cultured with growth factor supplementation remained viable. (**F**) Immunofluorescence of organoids co-cultured with GFP^+^ immortalized telocytes or supplemented with growth factors, stained for Ki67 or Sox9 on Day 4, shows that in co-cultured organoids, expression is restricted to the base of the budding region. In contrast, supplemented organoids exhibit broad expression along the entire budding region. Asterisk-staining in immortalized telocytes. Scale bar 100 µm.

We monitored organoid growth over four days. By day 2, organoids in all conditions - except those co-cultured with immortalized telocytes or supplemented with external niche factors - began to deteriorate (**Figures 5B-C** and **5E**). In contrast, organoids co-cultured with GFP^+^ immortalized telocytes not only maintained large sphere structures on days 1 and 2 but also began budding, differentiating, and forming crypt-like structures more prominently by day 3. Unlike tdTomato^+^ (telocyte-depleted) mesenchymal cells, which remained within the Matrigel, GFP^+^ immortalized telocytes extended and established close contact with the basal side of the epithelial organoids (**Figure 5D**), mimicking the positioning of telocytes in the intestine.

To evaluate stem cell activity, we compared organoids co-cultured with immortalized telocytes to those supplemented with external niche factors on day 4. Organoids were stained for Ki67, a proliferation marker and Sox9, a Wnt target gene expressed in intestinal stem and transit amplifying cells in vivo^49^ (**Figure 5F**). Interestingly, organoids co-cultured with immortalized telocytes showed restricted expression of Ki67 and Sox9 at the base of the budding epithelium, unlike the broader expression across the entire budding segment observed in organoids supplemented with external factors (+ENRF). This suggests that immortalized telocytes promote a more controlled stem cell activity and differentiation, more closely reflecting the in vivo environment. This underscores their significant potential as a key source of stem cell niche factors in vitro. Consequently, this cell line could be pivotal for investigating stem cell and niche interactions under both normal and pathological conditions.

### Immortalized telocytes support human organoid growth

To evaluate the clinical relevance of our immortalized telocyte cell line as a component of the stem cell niche, we tested its ability to support human organoids derived from normal colon samples without the addition of external niche factors. Remarkably, while human organoids cultured without external niche factors began to die by Day 2, those co-cultured with immortalized telocytes formed close associations with the telocytes and gradually developed cystic-like structures, similar to colon organoids grown in commercially enriched media (**Figures 6A-B**).

**Figure 6.**
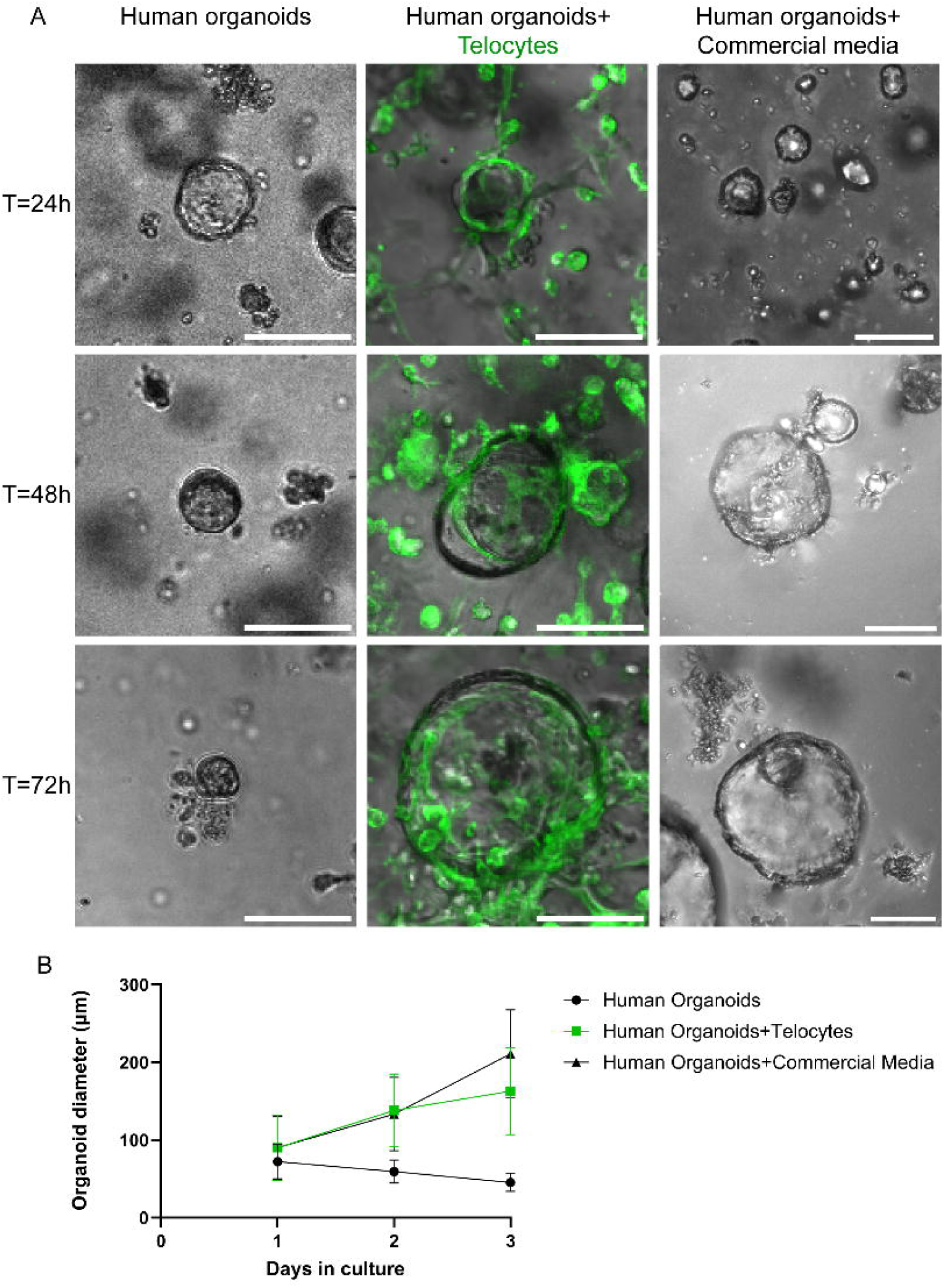
Immortalized telocytes support human organoid growth. (A) Photographs of human colon organoids show that organoids without GFP+ immortalized telocytes dye, whereas organoids co-cultured with GFP+ immortalized telocytes grow gradually similar to colon organoids cultured in enriched commercial media. Note the close contact between organoids and telocytes. (B) Quantification of organoid diameter reveals gradual growth in organoids co-cultured with GFP+ immortalized telocytes similar to organoids cultured in enriched media.

Notably, despite the telocyte cell line originating from the mouse small intestine and the organoids being derived from human colon tissue, the immortalized telocytes were still able to support organoid growth and preserve the original colon features. This suggests that telocytes may possess flexibility in their secretion profiles and niche-supporting factors, likely influenced by their interactions with the epithelium.

In conclusion, our findings highlight the potential of immortalized telocytes as a valuable resource for studying stem cell-niche interactions across both mouse and human models. This emphasizes the relevance of this cell line for applications in clinical research

## Discussion

Telocytes have recently been recognized as crucial components of the intestinal stem cell niche, yet their detailed biology and the specific mechanisms by which they sustain stem cell function and intestinal homeostasis remain largely unexplored. In this study, we developed an immortalized telocyte cell line that retains the cellular and molecular characteristics of native intestinal telocytes. These immortalized telocytes continue to express key niche factors and support the stem cell activity of both mouse- and human-derived organoids without the need for external supplementation.

Our research underscores the unique role of telocytes within the stem cell niche, as we demonstrated that mesenchyme lacking telocytes cannot support organoid growth. This finding suggests that immortalized telocytes retain critical stem cell niche functions in vitro, emphasizing their importance in maintaining stem cell niche activity.

Stem cells hold significant potential for applications ranging from regenerative medicine to patient-specific disease modeling, toxicology screening, and cultivated food production. However, directing stem cells to exhibit desired behaviors, such as expanding in a naïve state or differentiating into specific mature lineages, remains a challenge. The heterogenous networks of telocytes maintained in vitro, which express diverse signaling molecules that support both stem cell maintenance and differentiation, hold promise for more accurately recapitulating the natural differentiation process of stem cells. This approach provides a more native environment for controlling stem cell activity compared to the external addition of factor cocktails.

Co-culturing telocytes with organoids, rather than relying solely on niche factors, offers a valuable approach for studying the molecular and cellular mechanisms involved in stem cell growth and differentiation within their niche. This method enables the exploration of reciprocal communication and bidirectional signaling between stem cells and their niche-dynamic that cannot fully captured through external factors alone.

The establishment of an immortalized telocyte cell line will significantly enhance our understanding of basic telocyte biology, such as RNA localization along cellular extensions and the mechanisms they use to communicate with and signal to the epithelium. Advances in genome editing and CRISPR technology further enable the manipulation of both telocytes and organoids, offering new insights into stem cell-niche interactions in both homeostasis and disease.

Moreover, the ability of immortalized telocytes to support human organoid growth opens up opportunities to explore clinically relevant questions about telocyte interactions in both healthy and diseased states.

## Acknowledgments

We thank all members of M. Shoshkes laboratory for their helpful and insightful discussions. Special thanks to Dr. Yael Friedmann and the Bio-Imaging Facility at The Alexander Silberman Institute of Life Sciences, The Hebrew University, for their assistance with tissue processing and imaging using Transmission Electron Microscopy. This work was supported by a research grant from the Israel Science Foundation (grant #1997/19, M.S.-C). HUJI International PhD Talent Scholarship to A.G. Schematic illustrations were created with BioRender.com

## Author Contributions

M.C wrote the manuscript, conceived, carried out experiments, analyzed and interpreted the data. A.G carried out smFISH experiments and FACS analysis. D.D carried out Matrigel contraction assay and human organoid culture experiments; N.C carried out immunofluorescence staining. L.B.-S wrote the Helsinki protocol. Y.B-N, I.B-P, M.G, and M.S.-C designed and supervised the study. M.S.-C wrote the manuscript and directed the study.

## Methods

### Mouse models

Foxl1-Cre^50^ or Foxl1-CreERT2^51^ were crossed with Rosa-membrane-targeted dimer tomato protein (mT) or membrane targeted green fluorescent protein (mG) (Rosa-mTmG)^52^ (Jackson Laboratories, Bar Harbor, ME #007676).

All animal experiments were approved by the Animal Care and Use Committee of the Hebrew University of Jerusalem.

### Tamoxifen treatment

To induce Foxl1-CreERT2; Rosa-mTmG mice, tamoxifen (Sigma-Aldrich #10540-29-1) was dissolved in corn oil at a concentration of 30 mg/ml by shaking overnight at 37°C. The solution was then administered intraperitoneally at a dose of 150 mg/kg body weight. Tamoxifen was given over three consecutive days.

### Intestinal mesenchyme isolation and culture

Foxl1-Cre; Rosa-mTmG mouse jejunum was dissected and mesenchyme was isolated as previously described^13^. Briefly, the intestine was cut into 0.5 cm pieces. The epithelium was removed using 1 mM DL-dithiothreitol (DTT) and 2 mM EDTA. The remaining mesenchyme was then digested with 100 U/ml collagenase type VIII and 75 mg/ml DNase I in CM1640. Both primary and immortalized mesenchymal cells were cultured in DMEM medium containing 15% Fetal Bovine Serum and 1% Penicillin-Streptomycin.

### Immortalization by SV-40 Large T-antigen infection

FACS-sorted GFP^+^ telocytes and tdTomato^+^, mesenchymal telocyte-depleted cells were infected with viral particles expressing the SV40 Large T antigen (Addgene #170255), two weeks after sorting. Virus production followed standard procedures, involving transfection of a lentiviral vector encoding the protein, pHR-ΔR8.2 packaging vector, and the pCMV-VSV-G envelope plasmid into HEK293T cells. After transfection, supernatants containing viral particles were collected and concentrated by centrifugation. Intestinal mesenchymal cells were infected with 50μl of concentrated viral particles in 5-cm dishes, and selection was performed with Hygromycin (Thermo Fisher Scientific).

### FACS analysis

For single-cell suspension preparation, both primary and immortalized mesenchymal cells were incubated for 7 minutes with 0.25% Trypsin/1mM EDTA, then centrifuged at 1800 rpm for 5 minutes at 4°C. Single cells were resuspended in FACS buffer (5% FBS / PBS without Ca^2+^ and Mg^2+^). To optimize fluorescence compensation settings, anti-mouse (BD #552843) and anti-rat (BD #552845) beads were used. Cells were sorted based on GFP and tdTomato signals. For FACS analysis GFP^+^ immortalized Telocytes and tdTomato^+^ telocyte-depleted mesenchymal cells, were incubated for 30 minutes with the following antibodies:

| Antibody | Company | Catalogue number | Fluorophore |
| --- | --- | --- | --- |
| CD45 | BD | 563792 | BUV395 |
| CD31 | Biolegend | 102526 | PECF594 |
| GP38 | Biolegend | 127418 | APC-CY7 |
| CD34 | Biolegend | 119321 | BV421 |
| CD201 | Biolegend | 141506 | APC |
| CD90 | Biolegend | 140318 | BV605 |
| CD81 | Biolegend | 104912 | Percpcy5.5 |
| CD138 | Biolegend | 142514 | PE-CY7 |
| CD146 | Biolegend | 361022 | BV510 |
| CD321 | BD | 744657 | BV786 |
| CD249 | BD | 741743 | BUV737 |
| CD326 | Biolegend | 118245 | BV786 |

### Immunofluorescence

Cultured cells and organoids were fixed in 4% PFA for 20 minutes at room temperature. Fixed cells were incubated with primary and secondary antibodies diluted in CAS-Block (Invitrogen 008120) and imaged using a Nikon Eclipse Ti2 Confocal System (Nikon, Japan). Image processing was performed using Fiji or NIS Elements software packages. List of antibodies used in this study:

| Antibody | Company | Catalog number | Dilution |
| --- | --- | --- | --- |
| αSMA | Abcam | 5694 | 1:250 |
| αTubulin | Abcam | 7291 | 1:250 |
| CD34 | ThermoFisher | 14-0341-82 | 1:100 |
| CD81 | Abcam | 104912 | 1:100 |
| GFP | Abcam | 6673 | 1:400 |
| GFP | NovosBio | 100-1614 | 1:400 |
| Ki67 | Biolegend | 652402 | 1:100 |
| Mcam | Abcam | 34717 | 1:200 |
| MMP-10 | Abcam | 261733 | 1:100 |
| PDGFRα | R&D Systems | AF1062 | 1:100 |
| Porcn | ThermoFisher | PA5-43423 | 1:100 |
| Procr | Biolegend | 141505 | 1:100 |
| Sox9 | Millipore | AB5535 | 1:250 |
| Thy1 | Biolegend | 140318 | 1:100 |
| Vangl2 | Sigma-Aldrich | HPA025235 | 1:100 |

### Preparation of cells for Transmission Electron Microscopy imaging

Approximately 2,500 tdTomato^+^ telocyte-depleted mesenchymal cells and 750 GFP^+^ telocytes were cultured separately on an 8-chamber Lab-Tek slide and fixed in 2.5% glutaraldehyde and 2% paraformaldehyde prepared in 0.1M cacodylate buffer (pH 7.4) for 4 hours at room temperature, followed by overnight fixation at 4^0^C. The cells were rinsed four times for 10 minutes each in 0.1M cacodylate buffer, then post-fixed and stained with 1% osmium tetroxide and 1.5% potassium ferricyanide in 0.1M cacodylate buffer for 1 hour.

After staining, the cells were washed four times in cacodylate buffer and dehydrated through a graded ethanol series (30%, 50%, 70%, 80%, 90%, 95%, for 10 minutes each, followed by three washes in 100% anhydrous ethanol for 20 minutes each). Dehydrated cells were infiltrated with Agar 100 resin in ethanol at increasing concentrations (25%, 50%, 75%, and 100%) for 16 hours per step, then embedded in fresh resin and polymerized at 60^0^C for 48 hours.

The resin-embedded cells were sectioned into ultrathin slices (80 nm) using a diamond knife on Leica Reichert Ultracut S microtome. Sections were mounted onto 200-mesh-thin-bar copper grids and sequentially stained with uranyl acetate and lead citrate for 10 minutes each. The prepared sections were imaged using a Tecnai 12 TEM (120kV, Phillips, Eindhoven, the Netherlands) equipped with Phurona camera and RADIUS software (Emsis GmbH, Münster, Germany).

### RNA isolation and RT-qPCR

Passage 7 GFP^+^ immortalized telocytes and tdTomato^+^ mesenchymal telocyte-depleted cells were seeded on a 10cm Petri dishes and cultured. RNA was extracted using TRIzol reagent (Thermo Fisher,15596018) according to the manufacturer’s instructions. Briefly, approximately 5×10^6 cells per dish were incubated with 1ml of TRIzol. The cell lysate was then treated with chloroform and isopropanol to extract the RNA. RNA quantity and quality were assessed using NanoDrop™ and Qubit ™ (Thermo Fisher, A38189). Up to 1μg RNA was converted to cDNA using the iScript cDNA synthesis kit (Bio-Rad, 1725037), and qRT-PCR was performed in duplicate reactions using the LunaScript® RT SuperMix Kit (NEB, E3010). Expression values were normalized to HRPT-1 and GAPDH levels for GFP^+^ and tdTomato^+^ cells. mRNA levels were measured by TaqMan, and primer list is provided in the following table:

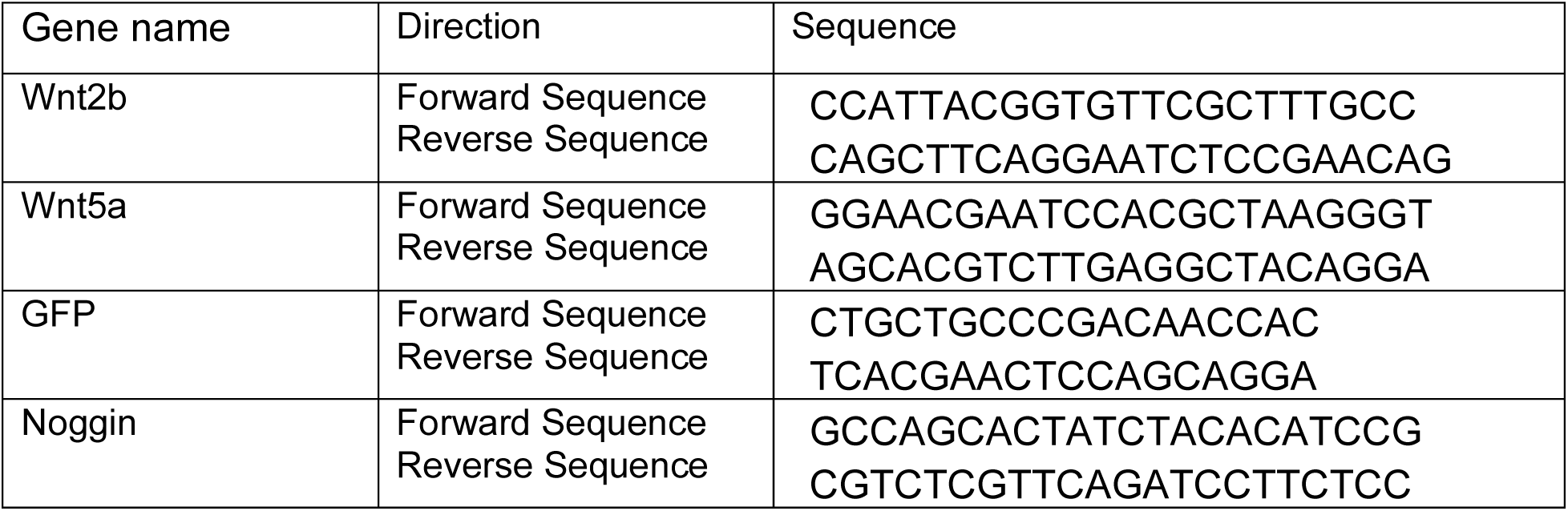
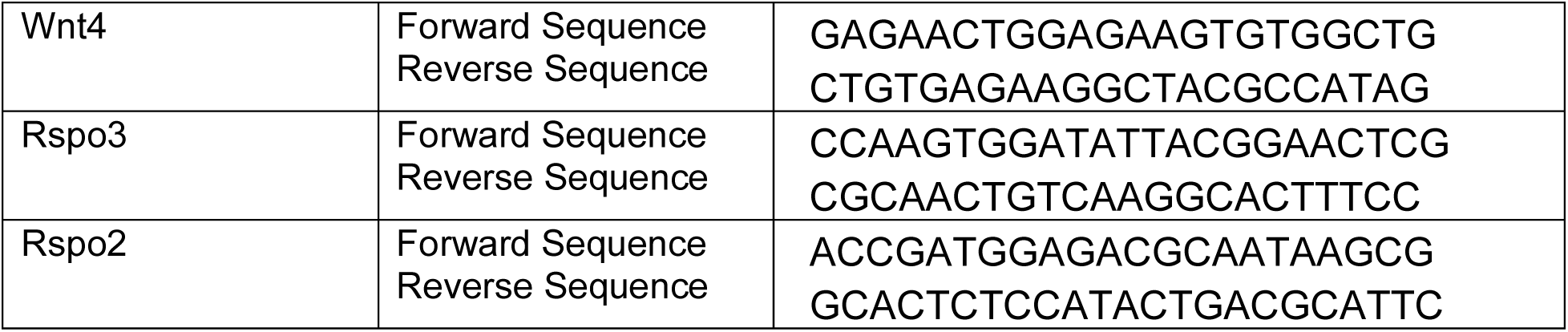

### Single-molecule RNA Fluorescence in-situ hybridization (smFISH)

Custom probe libraries, designed using the Stellaris FISH Probe Designer software (Biosearch Technologies, Inc., Petaluma, CA) and labeled with Cy5 or TMR were used to hybridize with the target coding RNA. For hybridization preparation, cells were seeded on coverslips and fixed with 3.7% formaldehyde for 10 minutes, following established protocol^53^. Antibody staining was performed by diluting the GFP antibody in hybridization buffer, followed by applying the Alexa 488 secondary antibody diluted in GLOX buffer for 20 minutes. Quantification of Foxl1 mRNA concentration was conducted using the TransQuant costum MATLAB program^54^.

### Matrigel contraction assay

GFP^+^ Immortalized telocytes were seeded at a density of 10,000 cells in 30µl Matrigel (BD Biosciences, 356230). Images of Matrigel domes were captured at 1,18, 42 and 92hours post-seeding. The circumference of Matrigel domes was measured using Fiji, and area quantification was analyzed and plotted in GraphPad Prism 9.5.1.

### Organoid-telocytes co-culture assay

Crypts were isolated from the jejunum of 10 to 15-week-old wild-type mice as previously described^55^. Briefly, the jejunum was flushed with ice-cold PBS, cut into pieces smaller than 5mm, and crypts were isolated by mechanical dissociation in PBS with 2mM EDTA on ice. Crypts were embedded onto Matrigel (BD Biosciences, 356230), 30μl per well in 24-well-plate. The culture medium was DMEM/F12 (Gibco) containing GlutaMax (1:100, Gibco) and Penicillin-Streptomycin (1:100 Biological Industries) supplemented with B-27 (1:50 Gibco), mouse Noggin (100 ng/ml Peprotech 250-38), mouse Rspo1 (500 ng/ml Peprotech, 315-32), mouse EGF (20 ng/ml Peprotech, 315-09), and human basic Fgf (10 ng/ml Peprotech, 450-33). The medium was changed every 2 days until co-culture.

To establish co-culture of murine organoids with GFP^+^ immortalized telocytes or tdTomato^+^ mesenchymal telocyte-depleted cells, organoids were dissociated, and mesenchymal cells were trypsinized (incubated 7 minutes in 0.25% Trypsin-1mM EDTA, then centrifuged at 1800 rpm for 5 minutes at 4°C). 10,000 mesenchymal cells were embedded onto 30μl Matrigel. The four different conditions were as follow: organoids, organoids+ supplements (ENRF), organoids+ immortalized telocytes, organoids+ immortalized mesenchymal telocyte-depleted cells. Images were acquired once a day for four consecutive days, and organoid diameters were measured using Fiji and analyzed with GraphPad Prism 9.5.1.

Human healthy colon organoids were obtained from the Organoid Center of Hadassah-Hebrew University Medical Organization, Jerusalem, under the Helsinki protocol HMO-20-921 and were co-cultured using the same protocol as for the mouse organoids.

Organoids, prior co-culture, were grown in Intesticult Organoid growth Medium (Human) from StemCell technologies.

## Supplemental Information

**Figure S1.**

**(A)** Immunofluorescence of immortalized telocytes stained for PDGFRα (red) showed all telocytes express PDGFRα and that two major subpopulations can be identified: PDGFRα^High^ and PDGFRα^Low^. (**B**) Immunofluorescence of immortalized telocytes stained for CD34 (red) shows a fraction of telocytes express CD34. (**C**) Immunofluorescence of immortalized telocytes co-stained for PDGFRα (red) and CD34 (white) shows co-expression of these markers in a subset of telocytes. (**D**) Immunofluorescence of immortalized telocytes stained for αSMA (red) and Thy1 (white), shows co-expression of αSMA and Thy1 in a subset of telocytes. Scale bar 10 µm.

**Video 1.**

Live imaging of GFP^+^ immortalized telocytes over 6 minutes (331 frames at 6 fps) reveals the dynamic structure of telocytes before and after cell division. The video highlights how telocytes establish and maintain interconnected networks. Scale bar 50 µm.

**Video 2.**

Live imaging of tdTomato^+^ immortalized (telocyte-depleted) mesenchymal cells over 92 minutes (573 frames at 12fps) highlights their size, morphology and reduced cell-cell interactions compared to telocytes.

