## Supplementary figures and images for "Immortalized Intestinal Telocytes - A Stem Cell Niche In Vitro"

### Supplementary S1

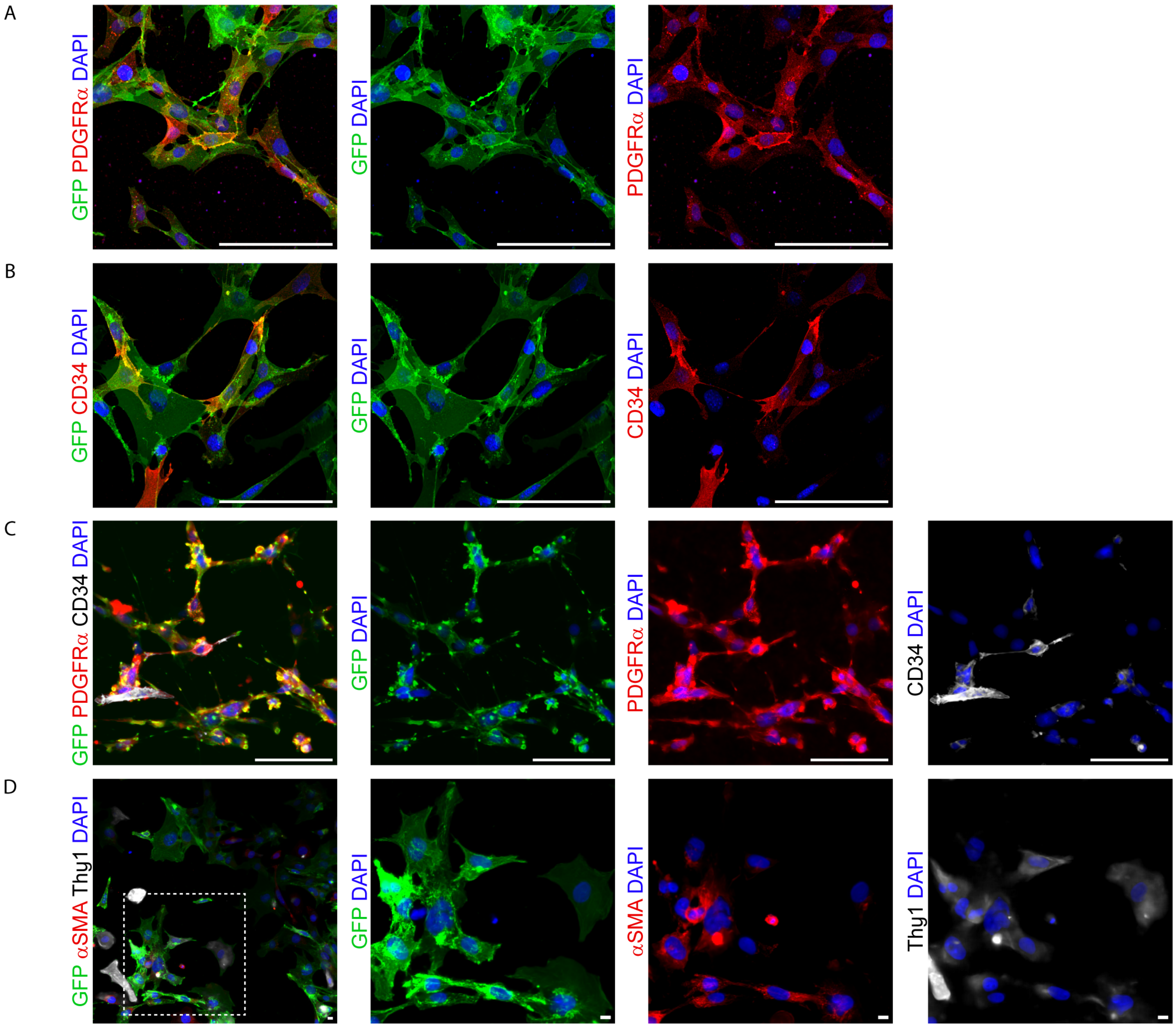
